# Cell-Type-Specific Expression of Renin-Angiotensin-System Components in the Human Body and Its Relevance to SARS-CoV-2 Infection

**DOI:** 10.1101/2020.04.11.034603

**Authors:** Hemant Suryawanshi, Pavel Morozov, Thangamani Muthukumar, Benjamin R. tenOever, Masashi Yamaji, Zev Williams, Thomas Tuschl

## Abstract

We have analyzed the cell-type-specific expression of the renin-angiotensin system (RAS) components across 141 cell types or subtypes as defined by single-cell RNA-seq (scRNA-seq) analysis. ACE2, one of the components of RAS, also facilitates SARS-CoV-2 entry into cells in cooperation with its associated protease TMPRSS2. Therefore, our analysis also contributes to the understanding of SARS-CoV-2 infection, spreading of the virus throughout the body, and potential viral interference with RAS in COVID-19 patients.

---

The COVID-19 pandemic caused by SARS coronavirus 2 (SARS-CoV-2) has created a global health emergency with more than a million confirmed cases as of April 4, 2020^1^. For cellular entry, SARS-CoV-2 relies on the interaction of its glycosylated viral spike (S) protein with the host membrane-bound aminopeptidase angiotensin-converting enzyme 2 (ACE2). Subsequently, transmembrane protease serine 2 (TMPRSS2) cleaves S and/or ACE2 protein, which facilitates the fusion of viral and cellular membranes^2^. ACE2 is also a crucial component of RAS, which regulates several key physiological processes such as blood pressure and electrolyte balance and viral infection of ACE2-expressing cells may impair RAS function^3^.

The secreted and systemically distributed protease renin (REN) activates RAS by proteolytically cleaving plasma angiotensinogen (AGT) to produce the 10-amino-acid (aa) hormonal peptide, angiotensin I (Ang I). Subsequently, Ang I is processed to the 8-aa Ang II by the metallopeptidase angiotensin-converting enzyme (ACE) present on the surface of pulmonary and kidney endothelial cells. Ang II is the active peptide in RAS triggering vasoconstriction, sodium retention, and fibrosis and signals by binding to its receptor AGTR1 or AT_1_R. ACE2, a metallopeptidase paralogous to ACE, counters the activity of ACE by digesting Ang II to the 7 aa Ang-(1-7) form, thereby attenuating the effects of Ang II^3^. Ang-(1-7) signals through binding to the G-protein coupled receptor MAS1 as well as the receptor AGTR2 or AT_2_R. Activation of MAS1 protein is coupled with several downstream pathways including activation of phospholipase A2 PLA2G4A, release of arachidonic acid, calcium-independent activation of nitric oxide synthase, activation of PI3K/Akt, MAP kinases, RhoA, and cAMP/PKA^4^.

Since SARS-CoV-2 interfaces with the RAS system by potentially competing for ACE2, we reviewed and analyzed 14 distinct tissue scRNA-seq datasets to determine cell-type- and tissue-specific expression of ACE2 and TMPRSS2 as well as other essential RAS factors including ACE, AGTR1, AGTR2, and MAS1 using both publicly available and unpublished datasets from our laboratory. The datasets include adult^5^ and fetal lung^6^, colon^7^, ileum^7^, esophagus^5^, rectum^7^, spleen^5^, fetal heart^8^, healthy and allograft kidney (data not published), skin^9^, first-trimester placenta and decidua^10^, and testis (data not published). In addition, we also analyzed bulk RNA-seq data of bronchoalveolar lavage fluid (BALF) from the first reported COVID-19 patient from Wuhan termed as ‘Wuhan BALF’ with no history of hepatitis, tuberculosis, or diabetes^11^, and normal human bronchial epithelial (NHBE) cells^12^, often used for studying coronavirus infection.

Based on published cell type assignment or by in-house generation of expression matrices using raw scRNA-seq datasets, we defined gene expression for 141 cell types including cell subtypes (Fig. 1). Epithelial cells across these different tissues showed the highest expression for the host factors ACE2 and TMPRSS2 required for viral entry^2^. In adult lung, alveolar type 1 and 2, as well as ciliated epithelial cells co-expressed ACE2 and TMPRSS2 with highest expression in alveolar type 2 cells. In the kidney, we found that proximal tubular cells and tubular progenitor cells co-expressed ACE2 and TMPRSS2. In the gastrointestinal tract, epithelial cells of the esophagus, rectum, colon, and ileum also co-expressed ACE2 and TMPRSS2. Furthermore, lung epithelial cell types additionally co-expressed ACE, AGTR1, and AGTR2, whereas kidney proximal tubular cells co-expressed ACE and AGTR1, but not AGTR2. Enterocytes of the gastrointestinal tract co-expressed ACE but only minimally either of AGTR1 or AGTR2. RAS is considered largely a ‘blood-borne hormonal’ system with endothelial cells being the major responders and this hypothesis is supported by the co-expression of ACE and AGTR1 in vascular endothelial cells in all tissues. However, AGTR2 expression was restricted to the vascular endothelial cells of lungs and fetal heart. ACE2, in contrast, was nearly absent from vascular endothelial cells of all adult tissues. The expression of the receptor MAS1 was tissue-specifically restricted to adult lung, esophagus, and epithelial progenitor cells of the gastrointestinal tract, but was absent from kidney.

**Fig. 1.**
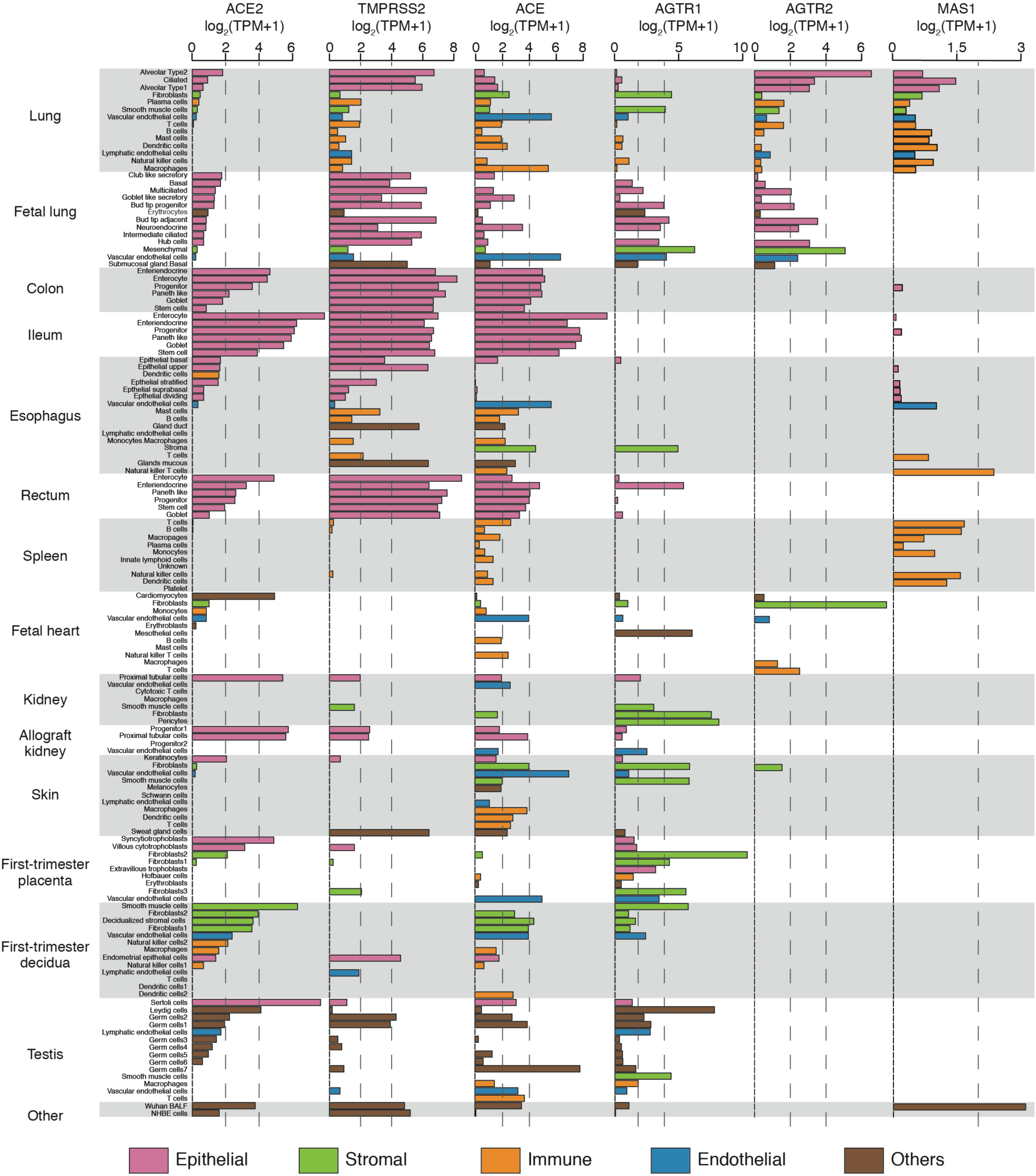
Gene expression distribution of the important components in SARS-CoV-2 infection and renin-angiotensin system (RAS) system across 141 cell types/subtypes from 14 tissues.

RAS undergoes major changes in response to pregnancy with the uteroplacental unit playing a crucial role in the signaling cascade^13^. The trophoblasts of first-trimester placenta (6-11 weeks of gestation) showed presence of ACE2 in syncytiotrophoblasts (SCTs) that form the outer layer of villous projections and the villous cytotrophoblasts (VCTs) located in the innermost chorionic villi layer while TMPRSS2 was restricted to the VCTs only. In addition, both SCTs and VCTs expressed AGTR1 but not ACE, AGTR2, or MAS1. Most strikingly, the first-trimester decidua, unlike other tissues, showed high ACE2 expression in stromal cells but lacked TMPRSS2 expression, which was largely restricted to the epithelial cells. Vascular endothelial cells of the first-trimester decidua, in addition to ACE2, expressed ACE and AGTR1 but not AGTR2, overall emphasizing the complex distribution of RAS components at the maternal-fetal interface.

The cell-type-resolved expression pattern of ACE2 suggests the possibility of direct involvement of organs other than airways and lung in SARS-CoV-2 infection. Acute gastrointestinal injury in critically ill patients and detection of SARS-CoV-2 RNA in stool samples^14^, points towards viral infection of epithelial cell types showing abundant expression of ACE2 and TMPRSS2 in colon, Ileum and rectum. Inflammation and myocardial injury is also associated with fatal outcome of COVID-19^15^. Expression of ACE2 in cardiomyocytes suggests the potential for direct infection of the virus in the heart. In the kidney, abundant ACE2 and TMPRSS2 co-expression in proximal tubular and progenitor cells also makes them potential targets of virus infection. Acute kidney injury (AKI) has been noted in a small but significant proportion of patients with COVID-19 disease and is an independent risk factor for in-hospital mortality^16^. Histopathology of kidney tissue obtained at autopsy in patients with SARS-CoV-2 infection showed severe acute tubular necrosis with lymphocytic infiltration and the presence of viral nucleocapsid proteins^17^. Several cell types in first-trimester placenta and decidua show abundant expression of host factors required for SARS-CoV-2 entry. This observation is crucial in the context of understanding whether the virus can be vertically transmitted from pregnant mother to the fetus. Such phenomenon remains controversial in absence of concrete evidence^18-20^. Co-expression of ACE2 and TMPRSS2 in Sertoli, Leydig and germ cells indicate potential pathogenicity to the testicular tissue. Interestingly, although viral RNA has been detected in several clinical specimens (termed ‘RNAaemia’ in absence of tests for presence of infectious viral particles) in the majority of infected patients, infectious virus particles have not been yet recovered from urine or blood of COVID-19 patients^21-24^. The detection of blood-circulating virus may be technically challenging, either because of its low titer in blood or because of recovering intact extracellular RNA from nuclease-rich biofluids^21^.

In summary, the unique cell-type specificity of RAS components revealed by scRNA-seq challenges certain aspects of the current paradigm of RAS^22^. The predominant epithelial cell distribution of ACE and ACE2 observed in multiple tissues is indicative of the presence of organ-centric RAS as opposed to the circulating RAS^23^. The discordance in the distribution of AGTR1 and AGTR2, as well as of MAS1, raises the intriguing possibility that additional receptors and downstream signaling pathways may be involved in RAS. Our finding of minimal expression of ACE2 in the vascular endothelial cells questions a role for Ang (1-7) as counterregulatory hormone of circulating RAS in attenuating the effects of Ang II^3^. Our analysis provides an important resource by highlighting cell types targetable by SARS-CoV-2 and towards understanding the tissue-wide expression of RAS components and its possible alterations in the context of COVID-19.

## Methods

### Source of scRNA-seq datasets and the downstream analysis

The first step in the analysis was to generate averaged expression profiles for the individual cell types using scRNA-seq datasets. For this purpose, the metadata containing cell type assignments to the barcodes and the raw UMI (unique molecular barcodes) count matrices were obtained for various tissues: ileum, colon, and rectum^7^, and fetal lung^6^. For other tissues such as adult lung, esophagus, and spleen published R object containing metadata and raw counts were obtained^5^. We recently published fetal heart^8^, skin^9^, and first-trimester placenta and decidua^10^ scRNA-seq data and it was used for generating averaged expression profiles for cell types. Next, we used in-house generated and unpublished scRNA-seq data from tissues of donor and allograft kidney, and testis. After averaged expression profiles for cell types were generated, the expression was normalized to a million to generate transcript per million (TPM) values, followed by transformation to log_2_(TPM+1) for representing the gene expression in Fig. 1.

## References

1. Coronavirus latest: confirmed cases cross the one-million mark. Nature (2020). doi:10.1038/d41586-020-00154-w

2. Hoffmann, M. et al. SARS-CoV-2 Cell Entry Depends on ACE2 and TMPRSS2 and Is Blocked by a Clinically Proven Protease Inhibitor. Cell (2020). doi:10.1016/j.cell.2020.02.052

3. Vaduganathan, M. et al. Renin-Angiotensin-Aldosterone System Inhibitors in Patients with Covid-19. N. Engl. J. Med. NEJMsr2005760 (2020). doi:10.1056/NEJMsr2005760

4. Karnik, S. S., Singh, K. D., Tirupula, K. & Unal, H. Significance of angiotensin 1-7 coupling with MAS1 receptor and other GPCRs to the renin-angiotensin system: IUPHAR Review 22. Br. J. Pharmacol. 17, 737–753 (2017).

5. Madissoon, E. et al. scRNA-seq assessment of the human lung, spleen, and esophagus tissue stability after cold preservation. Genome Biol. 21, 1–16 (2020).

6. Miller, A. J. et al. In Vitro and In Vivo Development of the Human Airway at Single-Cell Resolution. Dev. Cell 53, 117–128.e6 (2020).

7. Wang, Y. et al. Single-cell transcriptome analysis reveals differential nutrient absorption functions in human intestine. Journal of Experimental Medicine 217, 357 (2019).

8. Suryawanshi, H. et al. Cell atlas of the fetal human heart and implications for autoimmune-mediated congenital heart block. Cardiovasc. Res. 126, 1037 (2019).

9. He, H. et al. Single-cell transcriptome analysis of human skin identifies novel fibroblast subpopulation and enrichment of immune subsets in atopic dermatitis. Journal of Allergy and Clinical Immunology (2020). doi:10.1016/j.jaci.2020.01.042

10. Suryawanshi, H. et al. A single-cell survey of the human first-trimester placenta and decidua. Sci Adv 4, eaau4788 (2018).

11. Wu, F. et al. A new coronavirus associated with human respiratory disease in China. Nature 579, 265–269 (2020).

12. Gillen, A. E. et al. Molecular characterization of gene regulatory networks in primary human tracheal and bronchial epithelial cells. Journal of Cystic Fibrosis 17, 444–453 (2018).

13. Irani, R. A. & Xia, Y. The functional role of the renin-angiotensin system in pregnancy and preeclampsia. Placenta 29, 763–771 (2008).

14. Sun, J.-K. Acute gastrointestinal injury in critically ill patients with coronavirus disease 2019 in Wuhan, China. medRxiv 2020.03.25.20043570 (2020). doi:10.1101/2020.03.25.20043570

15. Guo, T. et al. Cardiovascular Implications of Fatal Outcomes of Patients With Coronavirus Disease 2019 (COVID-19). JAMA Cardiol (2020). doi:10.1001/jamacardio.2020.1017

16. Cheng, Y. et al. Kidney disease is associated with in-hospital death of patients with COVID-19. Kidney International (2020). doi:10.1016/j.kint.2020.03.005

17. Diao, B. et al. Human Kidney is a Target for Novel Severe Acute Respiratory Syndrome Coronavirus 2 (SARS-CoV-2) Infection. medrxiv.org

18. Zeng, L. et al. Neonatal Early-Onset Infection With SARS-CoV-2 in 33 Neonates Born to Mothers With COVID-19 in Wuhan, China. JAMA Pediatr (2020). doi:10.1001/jamapediatrics.2020.0878

19. Dong, L. et al. Possible Vertical Transmission of SARS-CoV-2 From an Infected Mother to Her Newborn. JAMA (2020). doi:10.1001/jama.2020.4621

20. Chen, H. et al. Clinical characteristics and intrauterine vertical transmission potential of COVID-19 infection in nine pregnant women: a retrospective review of medical records. The Lancet 395, 809–815 (2020).

21. Max, K. E. A. et al. Human plasma and serum extracellular small RNA reference profiles and their clinical utility. Proc. Natl. Acad. Sci. U.S.A. 115, E5334–E5343 (2018).

22. Pessôa, B. S. et al. Key developments in renin–angiotensin–aldosterone system inhibition. Nat Rev Nephrol 9, 26–36 (2013).

23. Campbell, D. J. Clinical relevance of local Renin Angiotensin systems. Front Endocrinol (Lausanne) 5, 113 (2014).

